# Visualizing *in vitro-in vivo* correlation of miktoarm copolymer nanomicelles in cancer cellular uptake and trafficking

**DOI:** 10.1101/755595

**Authors:** Zheng Cui, Xiaofei Zhang, Xiaojin Zhang, Suna He, Wei Gao, Bing He, Xueqing Wang, Hua Zhang, Zhenlin Zhong, Qiang Zhang

## Abstract

In this study, a clear correlation between the *in vitro and in vivo* cellular uptake and trafficking was discovered by delivering miktoarm copolymer nanomicelles (MCNs) to cancer cells and tumor tissues. To monitor this process, two different FRET pairs, DiO and DiI, DiD and DiR, were loaded into MCNs to monitor the Förster resonance energy transfer (FRET) efficiency. The change in FRET efficiency *in vitro* and *in vivo* demonstrated a similar sequence of events for the transport of MCNs: hyperbranched block PCL inserted into cytomembrane, while the loaded hydrophobic fluorescence probes were released and followed by time-dependent intracellular clustering within endocytic vesicles. Additionally, uptake of loaded fluorescence probes with successively increasing ratios of copolymers suggested that with the increase of mass ratio of copolymer to fluorescence probes, cellular uptake of probes significantly decreased. This result was also consistent with the uptake behavior in cancer tissues. Collectively, the interaction between MCNs and cellular membrane dictated the uptake and trafficking of core-loaded hydrophobic probes. This concept paves a new way to analyze *in vitro-in vivo* correlation of other nanocarriers for endocytosis mechanism studies as well as further novel copolymers design in biomedical applications.

---

Advances in nanotechnology have fostered to construct a diversity of nanomaterials with the ability to encapsulate and transport therapeutic or diagnostic agents^1-3^. Among these nanoscale delivery vehicles, polymeric micelles attracted the most attention in diverse biomedical fields as drug delivery vehicles^4-5^, contrast agents carriers^6-7^ and diagnostic devices^8-9^. The advantages of polymeric micelles used as drug delivery vectors include improving the solubility of poorly water soluble drugs with the amphiphilic structures^10^, stabilizing and protecting drugs that are sensitive to the surrounding environment^11^, reducing nonspecific uptake by the reticuloendothelial system (RES)^12^ and achieving the passive targeting delivery in solid tumors^13-14^. The conventional micelle drug delivery systems are mainly based on linear amphiphilic copolymers. However, linear amphiphilic copolymer micelles will dissemble *in vivo* when the concentration of the copolymer is diluted by the bloodstream to fall below the critical micelle concentration (CMC). As a result, the instability of linear amphiphilic copolymer micelles greatly limits their clinical application^15^.

To overcome the disadvantage of classical micelles, recent advances in synthetic methodologies have fostered to develop amphiphilic polymers with more complex architectures, including dendrimers, hyperbranched polymers, cyclic polymers, and star polymers^15-16^ in order to increase capacity of copolymer micelles and to enhance the stability of micelles *in vivo*. In this paper, we used novel amphiphilic miktoarm copolymers PEG_113_-(*hb*-PG)_15_-*g*-PCL_22_ with a linear-hyperbranched architecture bearing one monomethoxy poly(ethylene glycol) (mPEG) chain and several poly(ε-caprolactone) (PCL) chains on a hyperbranched polyglycerol (*hb*-PG) core^17^ (Figure 1A) to construct ultrastable miktoarm copolymer nanomicelles (MCNs). The highly hydrophobic branched structure of PCL chains offers inner porosity and imposes a robust steric barrier for embedded water insoluble drugs, which at the same time potentially allowing good release control. PEGylation is a common strategy to increase water solubility and compatibility in polymeric delivery systems. Moreover, densely PEG-coated MCNs enable their long circulation *in vivo* by evading the reticuloendothelial system (RES)^18^.

**Figure 1.**
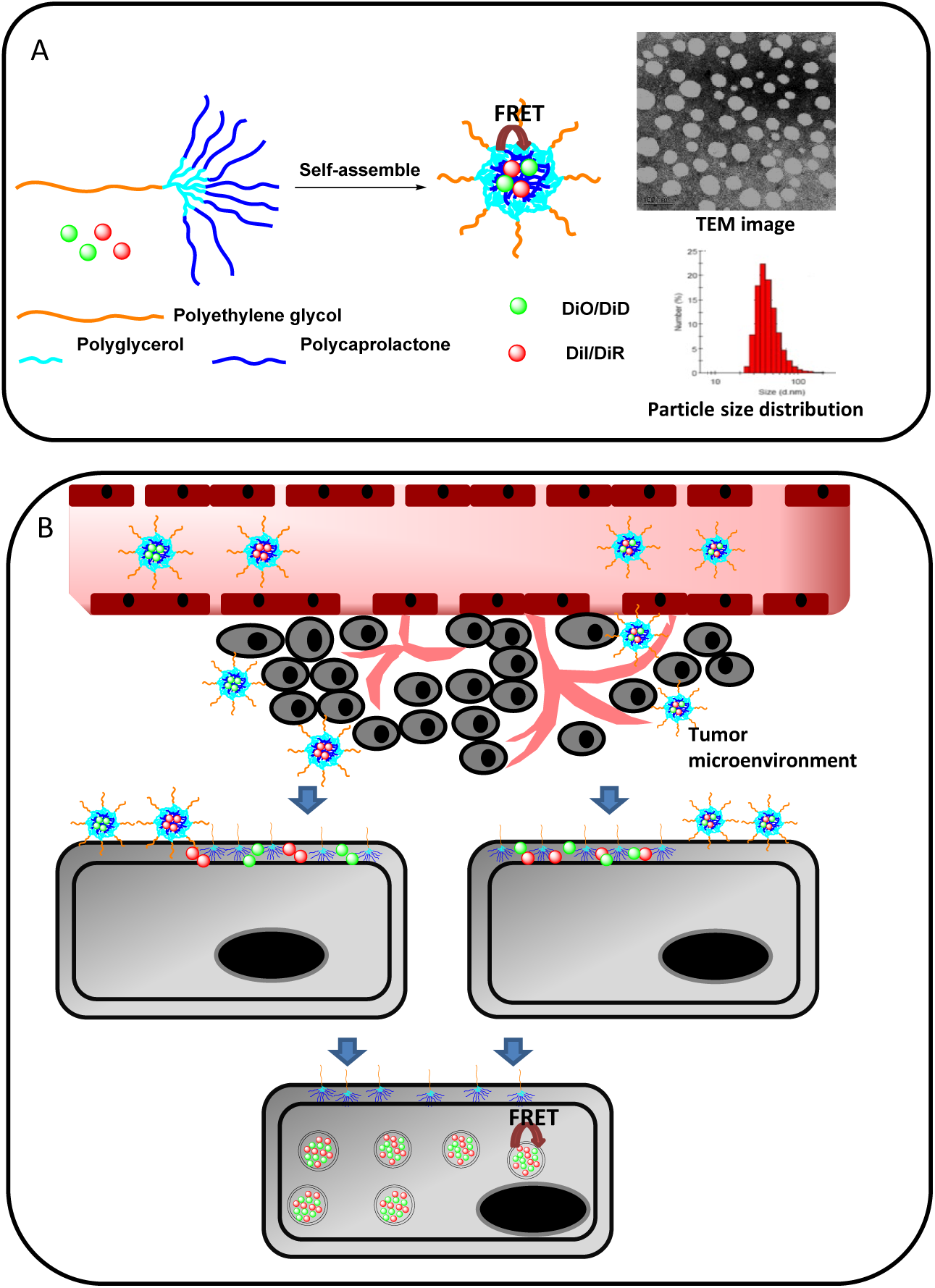
(A) Schematic illustration of miktoarm copolymer nanomicelles (MCNs), made of PEG_113_-(*hb*-PG)_15_-*g*-PCL_22._ (B) Schematic illustration of the concept of the study. Using the technique of Förster resonance energy transfer (FRET), the cellular uptake and trafficking behavior can be monitored.

Hitherto, many drug-incorporated polymeric micelles are currently undergoing clinical trials^19-21^, which contributes to our understanding of the pathways involved in polymeric micelle delivery^22^. However, to our knowledge, no studies have examined the cellular uptake mechanism of miktoarm copolymer micelles. Furthermore, despite a great deal of insight into the endocytosis mechanism of classical micelles *in vitro*^23-24^, *in vitro-in vivo* correlation of nanomicelle transport is still unknown because no nanoscale direct visualization of how the nanomicelles are internalized and transported have been shown. Most current studies of intracellular trafficking of nanomicelles are mainly limited to co- localization of nanomaterials with specific endocytic markers or the exclusion of specific mechanisms by chemical inhibition or cell mutation^25-26^.

In an attempt to address these problems mentioned above, we investigated cancer cellular uptake and trafficking correlation of PEG113-(*hb*-PG)15-*g*-PCL22 MCNs *in vitro* and *in vivo*. The basic concept of the entire study was illustrated in Figure 1B. The current work characterized the physical properties of MCNs, tested their internalization modalities of dual-labeled MCNs, visualized the release and transport process with *in vitro* and *in vivo* FRET imaging, and began to determine their underlying internalization mechanisms employing successively increasing ratio of copolymers. We show here that cellular uptake of loaded hydrophobic probes is much faster than labeled copolymers. The transport process *in vitro* and *in vivo* involved a clear sequence of events: hyperbranched block PCL inserted into the cytomembrane, while the loaded hydrophobic fluorescence probes released, followed by time-dependent intracellular clustering within endocytic vesicles. With the increase of mass ratio of copolymer, the uptake amount and speed of loaded fluorescent probes significantly decrease, which resulted in slower *in vivo* clearance rates of loaded fluorescent probes in the peripheral region of the tumor.

The reason for choosing these four kinds of fluorescent probes (DiO, DiI, DiD and DiR) in this study was as follows. (1) DiO, DiI, DiD and DiR are all hydrophobic fluorescence probes and thus can be used to represent hydrophobic drugs loaded in MCNs. (2) The fluorescent probes, DiO and DiI, are used in cellular assays, while the near-infrared fluorescent probes DiD and DiR-loaded MCNs are used in animal studies. In this case, the similar chemical structures of DiO, DiI, DiD and DiR could aid us to construct dye-loaded MCNs with similar release behavior in order to discover the *in vitro -in vivo* correlation. (3) The FRET pair DiO and DiI, with DiO as donor and DiI as acceptor, was used to monitor FRET efficiency *in vitro*; while the second FRET pair DiD and DiR, with DiD as donor and DiR as acceptor was used to detect the change of FRET efficiency *in vivo*.

## Materials and methods

### Materials

PEG_113_-(*hb*-PG)_15_-*g*-PCL_22_ (Mn=8500) was kindly donated by Dr. Zhenlin Zhong (Wuhan University)^17^. Sulforhodamine B (SRB), trichloroacetic acid (TCA) and Tris base were all purchased from Sigma-Aldrich (St. Louis, MO, USA). Hoechst 33258 was purchased from Molecular Probes, Inc (Oregon, USA). DiO and DiI were provided by Beyotime Institute of Biotechnology (Shanghai, China); DiD and DiR were purchased from Biotium, Inc (Hayward, USA).

### Preparation of DiO or/and DiI, DiD or/and DiR-loaded MCNs

DiO and DiI, DiD and DiR were chosen as FRET pairs for the following study. All the micelles were prepared using thin film hydration method^27^. Briefly, dyes (DiO, DiI, DiD, DiR), PEG-(*hb*-PG)-*g*- PCL were codissolved in acetonitrile. The acetonitrile was evaporated under vacuum with a rotary evaporator at 40 °C. The obtained copolymer film was hydrated in RPMI- 1640 medium, followed by stirring in the 40 °C water bath for 5 min. Finally, the solution was filtered through a 0.22 μm membrane. The mass ratio of dye to copolymer was 1:500, 1:750, 1:1500 and 1:3000. The final concentration of DiO or DiI for the cell uptake assay was 2 μg/mL.

### Characterization of DiO- or/and DiI-, DiD- or/and DiR-loaded MCNs

The particle size and zeta potential of the prepared blank MCNs and dye-loaded MCNs were measured by dynamic light scattering (DLS) using Malvern Zetasizer Nano ZS (Malvern, UK) at 25 °C^28^. The morphology of blank MCNs after dilution was investigated by transmission electron microscope (TEM, JEOL, Japan) after negative staining with uranyl acetate solution (1%, w/v). Encapsulation efficieny (E.E) was defined as the actual amount of fluorescent probes encapsulated as detected by the fluorospectrophotometer divided by the original amount of fluorescent probes. The probe-loaded micelles were diluted by acetonitrile and the concentration of fluorescence probes was measured using fluorospectrophotometer^29^.

The critical micelle concentration (CMC) was determined using pyrene as a fluorescent probe^4^. The concentration of block copolymer varied from 0.05 μg/mL to 50 μg/mL and the concentration of pyrene was fixed at 6×10^−6^ M. The fluorescence spectra were recorded using a Cary Eclipse fluorescence spectrometer using an emission wavelength of 390 nm. The excitation spectra were recorded ranging from 300 nm to 360 nm. The emission fluorescence at 333 nm and 335 nm was monitored. The CMC was estimated as the cross point when extrapolating the intensity ratio I_335_/I_333_ at low and high concentration regions.

### Cell Culture

Human breast cancer MCF-7 cells were obtained from the Institute of Basic Medical Science, Chinese Academy of Medical Sciences (Beijing, China). Cells were cultured in RPMI-1640 medium (M&C Gene Technology, Beijing, China) supplemented with 10% fetal bovine serum (FBS), 100 units/mL penicillin and 100 μg/mL streptomycin at 37 °C in a humidified atmosphere containing 5% CO_2_. The cells for all experiments were in the logarithmic phase of growth^30^.

### Cytotoxicity assay

The cytotoxicity of MCNs was evaluated *in vitro* using the SRB (sulforhodamine B) and LDH (lactate dehydrogenase) assays. SRB assay was performed on MCF-7 cells as originally described. Briefly, MCF-7 cells were seeded into 96-well culture plates at 3000-3500 cells/well and grown at 37 °C in the presence of 5% CO_2_ for 24 h. The final concentration of MCNs was in the range of 0.025-6 mg/mL. The culture medium without any MCNs was used as a blank control. 9% Triton X-100 was added as a positive control. After exposure for 24 h, the cells were fixed with 10% trichloracetic acid (TCA) at 4 °C for 1 h, the 96-well plate was then washed 4 times with water, while vigorously flicking the plate between washes to remove excess water. SRB solution was added into each well and incubated for 30 mins, after which the excess dye was removed by washing repeatedly with 1% acetic acid. After air-drying, the bound dye was re-dissolved in 10 mM Tris base solution for OD determination at 565 nm using a microplate reader (Bio-Rad 680, America).

As an additional test, we also measured the leakage of LDH in the culture medium. The assay relies on measuring the activity of LDH in catalyzing the reaction: 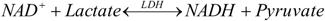. The culture medium without any MCNs was used as a blank control. 9% Triton X-100 was added as a positive control. After treatment with MCNs (6 mg/mL, 1 mg/mL) for 24 h, media were collected and centrifuged, and the activity of LDH released from the cytosol of damaged cells was assessed using the LDH Cytotoxicity Assay Kit (Applygen Technologies Inc, China). The maximum releasable LDH activity in the cells, induced by the addition of 9% Triton X- 100, was measured and used as 100% LDH released. The optical density at a wavelength of 440 nm was measured using a microplate reader (Bio-Rad 680, America). The formula below was used to calculate LDH release percentage.

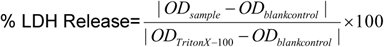

### Flow cytometry analysis

Approximately 6.0×10^5^ MCF-7 cells/well were seeded in 6-well plates and cultured for 24 h at 37 °C. Cells were exposed to various DiO and DiI co-loaded MCNs (the mass ratio of dyes and copolymers were 0, 1:500, 1:750, 1:1500 and 1:3000, respectively) at 37 °C for 1 h (or longer periods as indicated in the legend) for cellular internalization studies. Concentration of DiO and DiI in all formulations was 2 μg/mL. Cells were then washed and trypsinized, centrifuged, resuspended in pre-cooled PBS. Flow cytometry was performed on a FACScan flow cytometer (FACScan, Becton Dickinson, San Jose, CA).

### Spectroscopic characterization of dyes

The emission spectra were recorded on a Cary Eclipse ultraviolet/visible light/near-infrared spectrophotometer (Varian Inc.) using 1-cm quartz cells. Samples included DiO-loaded MCNs (DiO MCNs), DiI-loaded MCNs (DiI MCNs), DiO- and DiI-co-loaded MCNs (DiO/DiI MCNs), DiO-loaded MCNs mixed with DiI-loaded MCNs (DiO MCNs & DiI MCNs Mixtures), and DiO/DiI MCNs destroyed by 10×acetonitrile. The samples above were excited at 484 nm, and the emission spectra of the donor/acceptor were recorded at 490-600 nm for FRET measurements. Moreover, samples including DiD-loaded MCNs (DiD MCNs), DiR-loaded MCNs (DiR MCNs), DiD- and DiR-coloaded MCNs (DiD/DiR MCNs), and DiD-loaded MCNs mixed with DiR-loaded MCNs (DiD MCNs & DiR MCNs Mixtures) were excited at 644 nm and the emission spectra of the donor/acceptor recorded at 650-800 nm for FRET measurements.

### Intracellular monitoring fate of hydrophobic probes using FRET

To monitor the fate of hydrophobic dyes released from MCNs, MCF-7 cells were treated with DiO MCNs, DiI MCNs, DiO/ DiI MCNs and DiO MCNs & DiI MCNs Mixtures in the serum-free 1640 medium, respectively. After incubation at predetermined time intervals, cells were washed twice with pre-cooled PBS before they were fixed in 4% paraformaldehyde. The fixed cells were visualized and examined using a Leica TCS SP5 confocal microscope. Confocal images were acquired with the excitation at 488 nm. The emission wavelength was between 555-655 nm for DiI (Acceptor) detection and the emission wavelength was between 500-530 nm for DiO (Donor) detection.

### Tumor Implantation

Female BALB/c nude mice (18-20 g) were purchased from Vital Laboratory Animal Center (Beijing, China) and acclimated at 25°C and 55% humidity under natural light/dark conditions for 1 week before the study, with free access to standard food and water (Vital Laboratory Animal Center, Beijing, China). MCF-7 tumor-bearing mice were prepared by inoculating 4×10^6^ MCF-7 cells in the right flank of female BALB/c nude mice. When tumor volume reached about 300 mm^3^, MCF-7 tumor bearing mice were randomly assigned to groups for live imaging studies. All procedures were approved by the Institutional Animal Care and Use Committee at Peking University Health Science Center and were in accordance with international guidelines on the ethical use of animals^31-32^.

### *In vivo* monitoring the uptake and trafficking of hydrophobic NIRF FRET probes

To verify the role of MCNs in delivering hydrophobic molecules to xenografted MCF-7 tumors, hydrophobic NIRF FRET probes (DiD, DiR) were utilized. Animal model of female BALB/c mice xenografted MCF-7 tumors were established in the same way as discussed above. Animals in two groups received the following two treatments respectively: DiD/DiR MCNs and DiD MCNs & DiR MCNs Mixtures. The dose of DiD and DiR in this study was 100 μg/kg. Near-infrared fluorescence(NIRF) imaging experiments were performed using a Kodak multimodel imaging system (Carestream Health, Inc. USA) at 1 h, 3 h, 5 h, 8 h, 12 h and 24 h. FRET images were acquired with an excitation bandpass filter at 610 nm and an emission bandpass filter at 790 nm. DiD images were acquired with an excitation bandpass filter at 610 nm and an emission bandpass filter at 720 nm. DiR images were acquired with an excitation bandpass filter at 720 nm and an emission bandpass filter at 790 nm. Fluorescence exposure time was 2 min and X-Ray exposure time was 30 s per image. All images were normalized and analyzed using the Carestream MI SE software.

### *In vivo* distribution studies by live imaging studies

In order to observe the real-time distribution and tumor accumulation ability of various fluorescence DiR MCNs formulations in PBS, including the mass ratio of DiR to copolymers of 1:500, 1:750, 1:1500 and 1:3000, non-invasive *in vivo* imaging systems were utilized on the female BALB/c mice xenografted MCF-7 tumors. The mice were injected with 0.2 mL of various DiR MCNs listed above (100 μg/kg DiR) respectively, via the tail vein, and then anesthetized by 2% isoflurane delivered via a nose cone system before scanning. Near-infrared fluorescence (NIRF) imaging experiments using a Kodak multimodel imaging system (Carestream Health, Inc. USA) at 5 h, 8 h, 12 h, 24 h, 48 h, 60 h, 72 h and 96 h with an excitation bandpass filter at 720 nm and an emission bandpass filter at 790 nm. Fluorescence exposure time was 2 min and X-Ray exposure time was 30 s per image.

### Intratumoral distribution of hydrophobic probes

To observe dye uptake and distribution into MCF-7 tumors, female BALB/c mice xenografted MCF-7 tumors were administered DiO/ DiI MCNs, DiO MCNs and DiI MCNs Mixtures in PBS via injection into the tail vein (100 μg/kg DiR). The mass ratio of DiO and DiI to copolymers was 1:750. The concentration of DiD and DiR was 100 μg/kg. After 12 h, the mice were sacrificed, and tumor tissues were harvested, placed in Tissue-Tek OCT embedding medium (Sakura Finetek, Tokyo, Japan), frozen on dry ice, sectioned using a cryostat, mounted on slides, and then fixed in 4% paraformaldehyde. The sections were incubated with Hoechst 33258 (1μg/mL)^33^ for 30 min at room temperature followed by washing with PBS. The stained tumor cryosections were examined using CLSM (Leica TCS SP5).

## Results and discussion

### Characterization of MCNs

Miktoarm copolymers have received increased attention because of their stability in forming nanomicelles and flexibility in their synthesis for biomedical applications. In this study, we used amphiphilic miktoarm copolymers PEG_113_-(*hb*-PG)_15_-*g*-PCL_22_ to construct nanomicelles. Figure 1 shows the schematic illustration of self-assembled MCNs. The hydrophobic PCL segments are locked in the dense inner core of micelles, while the hydrophilic PEG and PG chains formed the corona shell. MCNs encapsulated with five different combinations of hydrophobic fluorescence probes were prepared and characterized. The hydrodynamic diameters of the five MCNs with different encapsulations were approximately 50 nm with polydispersity indices (PDI) well below 0.22 (Table S1, Figure 1). The zeta potential of five MCNs was all around 0 mV, which indicated that encapsulation did not affect the zeta potential of MCNs. The morphology of spherical DiO and DiI co-loaded with MCNs (DiO/DiI MCNs) was observed in the TEM image (Figure 1). All encapsulation efficiencies of the five MCNs were around 90% (Table S1), meaning that MCNs have great potential in loading water-insoluble drugs if used to construct drug delivery systems. According to the excitation spectra of pyrene at different polymer concentrations, the CMC of PEG_113_-(*hb*-PG)_15_-*g*-PCL_22_ miktoarm copolymer nanomicelles was determined to be 0.37 µg/mL.

### Cytotoxicity test for MCNs

Cytotoxicity is a main concern in the application of drug delivery systems. The commonly used MCF-7 breast cancer cells were chosen for this study and the following *in vivo* assays. To examine the cytotoxicity of MCNs, we incubated MCF-7 cells with MCNs for 24 h. The toxicity of MCNs to MCF-7 cells was measured by the SRB assay^34^. Beyond our expectations, MCNs increased cell viability slightly in all groups as shown in Figure S1A. No statistically significant differences in toxicity were observed between the groups treated with different concentrations of MCNs samples. The LDH assay is a method of measuring the membrane integrity as a function of the amount of cytoplasmic LDH leaked into the medium^35^. To test membrane rupture, LDH assay was utilized on MCF-7 cells. 1 mg/mL and of 6 mg/mL MCNs used in our following cellular assay was tested in the LDH assay. The LDH leakage in the medium for the two groups did not significantly increase (P>0.05, Figure S1B). These results indicated that there was no significant cytotoxicity observed for the MCNs on MCF-7 cells.

### Intracellular trafficking studies of FRET MCNs on MCF-7 cells. *A) FRET measurements*

To monitor the release process of core-loaded hydrophobic molecules from MCNs *in vitro*, FRET pair DiO (Donor) and DiI (Acceptor) co-loaded MCNs (DiO/DiI MCNs) were prepared. The FRET of DiO/DiI MCNs was investigated with a fluorospectrophotometer. Figure 2A shows the fluorospectra of MCNs loaded with DiO (DiO MCNs), MCNs loaded with DiI (DiI MCNs), DiO/DiI MCNs, mixtures of MCNs loaded with DiO and MCNs loaded with DiI (DiO MCNs & DiI MCNs Mixtures). Energy transfer was evaluated by the intensity ratio of acceptor to donor (*I*_DiI_/*I*_DiO_), calculating the intensities of the acceptor emission (565 nm) and the intensities of the donor emission (501 nm) using the excitation at 484 nm. For DiO/DiI MCNs, DiO and DiI was condensed together by MCNs, with a FRET ratio of 2.40. When DiO-loaded MCNs and DiI-loaded MCNs were prepared respectively and mixed with each other (DiO MCNs & DiI MCNs Mixtures), no FRET existed with a FRET ratio of 0.25. DiO and DiI were not directly condensed with each other, thus FRET could not occur. After DiO/DiI-loaded MCNs decomposed by acetonitrile, the FRET signal disappeared because DiO and DiI could not be closely condensed any more, resulting in a FRET ratio of 0.08. Figure 2B shows the CLSM image of DiO/DiI loaded MCNs and its decomposition by acetonitrile. For DiO/DiI MCNs, the fluorescence intensity of the donor DiO strongly decreased in the presence of the acceptor DiI, and simultaneously, the emission of DiI was enhanced. When decomposed by acetonitrile, the fluorescence intensity of the donor DiO recovered. At the same time, FRET-mediated DiI emission strongly decreased.

**Figure 2.**
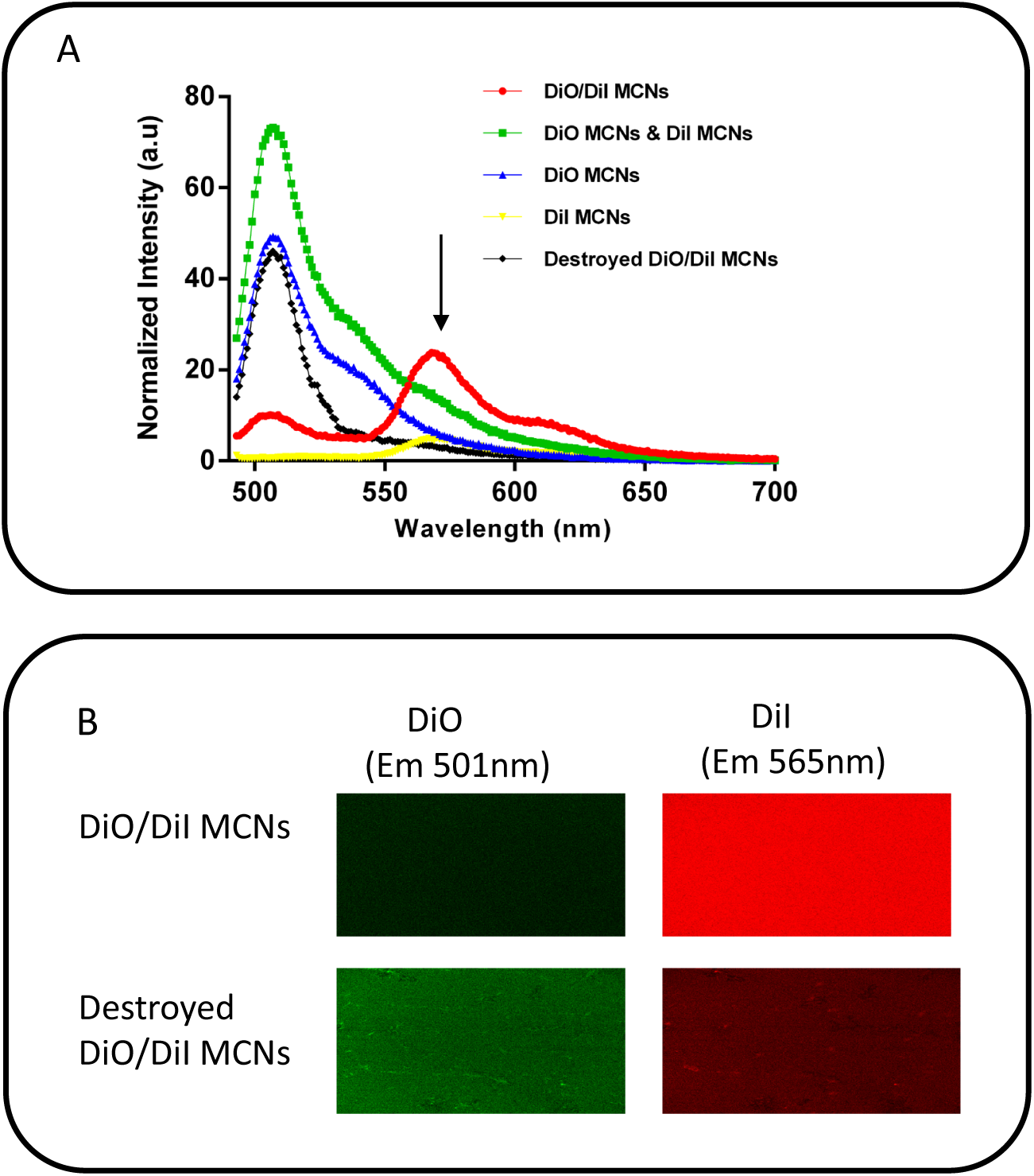
Fluorospectra and confocal laser scanning microscopy (CLSM) image of MCNs. (A) Fluorospectra of MCNs loaded with DiO (DiO MCNs), MCNs loaded with DiI (DiI MCNs), MCNs co-loaded with DiO and DiI (DiO/DiI MCNs), mixtures of MCNs loaded with DiO and MCNs loaded with DiI (DiO MCNs & DiI MCNs) and DiO/DiI MCNs decomposed by acetonitrile (Destroyed DiO/DiI MCNs). (B) CLSM image of DiO/DiI MCNs and acetonitrile destroyed DiO/DiI MCNs. Green fluorescence represents the fluorescence of DiO. Red fluorescence represents the fluorescence of DiI. The concentration of DiO was 2 μg/mL, the concentration of DiI was 2 μg/mL for all formulations that has DiO or DiI.

### B) FRET efficiency comparison between FRET Sensitized Emission (SE) method and FRET Acceptor Photobleaching (AB) method

A proper method to analyze the FRET efficiency in MCF-7 cells was required to evaluate the endocytosis behavior of MCNs. Sensitized Emission (SE) is one of the most popular methods for evaluation of FRET efficiencies. This method involves measuring the donor and the FRET signal (donor excitation only) in sequence with the detection of the acceptor (acceptor excitation only)^36^. FRET Acceptor Bleaching is another frequently used method for FRET efficiency measurements *in vitro*. The method involves measuring the donor “de-quenching” in the presence of an acceptor. This can be done by comparing donor fluorescence intensity in the same sample before and after destroying the acceptor by photobleaching. If FRET was initially present, a resultant increase in donor fluorescence will occur after photobleaching the acceptor^37^. Both methods were used to evaluate FRET efficiency in MCF-7 cells 4h post-DiO-DiI MCNs incubation. Five circled regions were selected accordingly to measure the FRET efficiency by both the FRET SE method (Figure S2A) and FRET AB method (Figure S2B). After t- test calculation with a significance level under 5%, there was not enough evidence to show that data obtained from the two methods were different. Considering that the FRET AB method required 8 s to photobleach each region before measuring FRET efficiencies, the FRET SE method is less time-consuming if dealing with the same amount FRET samples. Moreover, it is possible to bleach other regions accidentally if frequently photobleaching each spot and introduce error in the FRET efficiency measurement. As a result, the FRET SE method was utilized to monitor the dynamics of cellular endocytosis in the following experiment.

### C) Intracellular fate of FRET MCNs in MCF-7 cells

We continued to investigate the intracellular fate of two kinds of MCNs (DiO/DiI MCNs and DiO MCN & DiI MCN Mixtures) in MCF-7 cells using FRET imaging. CLSM images showed the representative intracellular fate of DiO/DiI MCNs and DiO MCNs & DiI MCNs Mixtures in the majority of the cells (Figure 3). The FRET SE method was used to quantify FRET efficiencies by the spectra intensities recorded from 30 fluorescent areas inside the cells. By incubating MCF-7 cells with DiO/DiI MCNs at 37 °C for 1 h, we observed the significantly diminished FRET effect (Figure 3A). The presence of co-localization of DiO and DiI but absence of FRET fluorescence from the plasma membrane might suggest a release of the two fluorescent probes from MCNs during cellular endocytosis of MCNs. It is also possible that not enough micelles were uptaken by cells within the short incubation time. At 2 h and 4 h incubation, the fluorescence signals of FRET-mediated DiI became stronger. The results indicated that DiO and DiI were likely released from MCNs and concentrated in endocytic vesicles in the process of cellular endocytosis and intracellular sorting, which led to the recovery of FRET signals. More interesting results were observed from the group of DiO MCNs & DiIs MCNs Mixtures (Figure 3B). DiO MCNs & DiI MCNs Mixtures had no FRET signals because of separate encapsulation of the two fluorescence probes. However, after incubating MCF-7 cells with DiO MCNs & DiI MCNs Mixtures for 2 h and 4 h at 37 °C, the FRET efficiencies were almost the same with that of the DiO/DiI MCNs’ (Figure 3C). The results from both groups further proved that DiO/DiI MCNs and DiO MCNs & DiI MCNs Mixtures have similar pathways in cellular trafficking.

**Figure 3.**
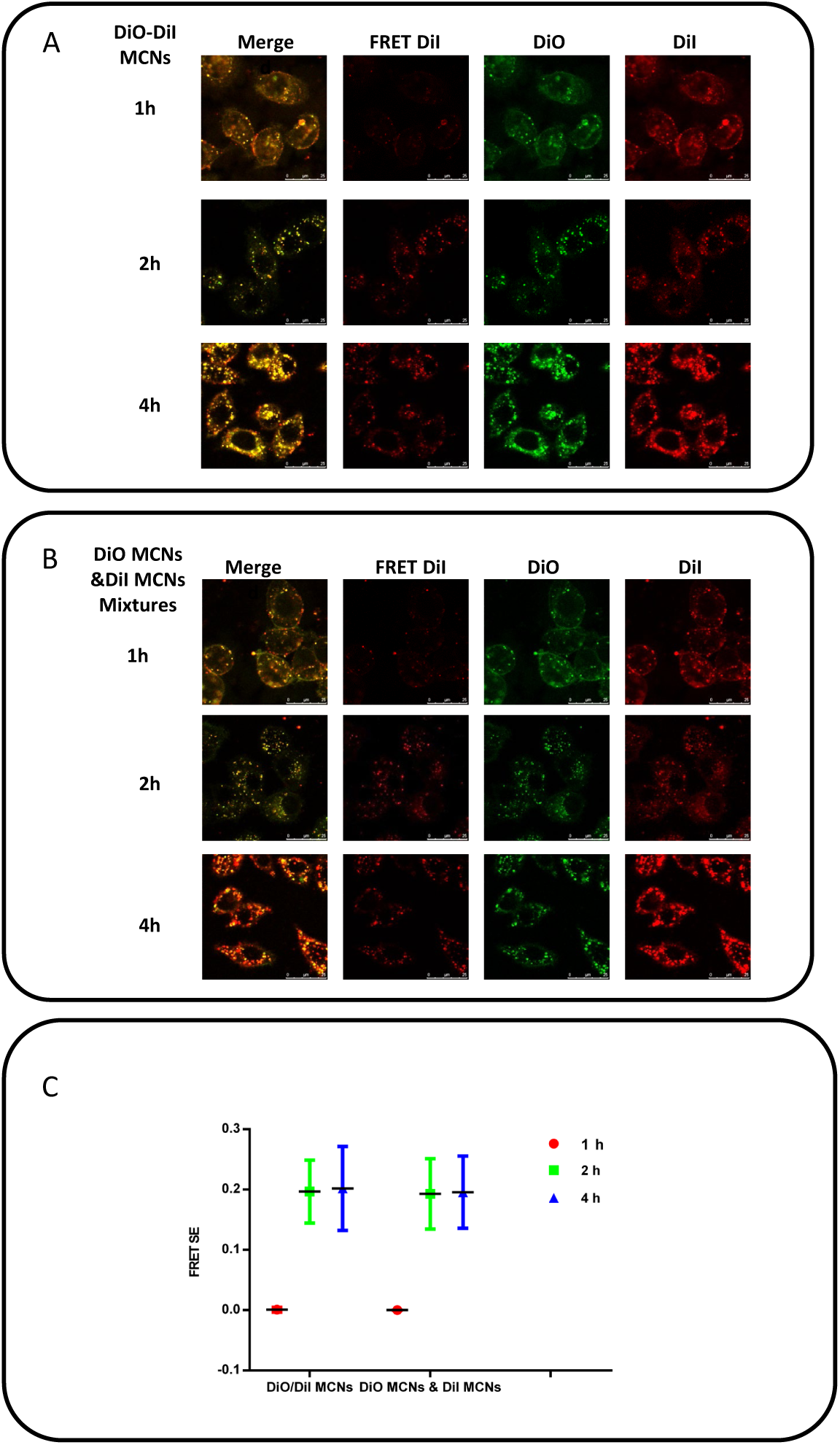
Intracellular trafficking studies of DiO/DiI MCNs and DiO MCNs & DiI MCNs Mixtures to MCF-7 cells via CLSM. (A) CLSM images of MCF-7 cells incubated with 1.5 mg/mL DiO/DiI MCNs (loaded with 0.16% DiO and 0.11% DiI). (B) CLSM images of MCF-7 cells incubated with 1.5 mg/mL DiO MCNs & DiI MCNs Mixtures (loaded with 0.16% DiO and 0.11% DiI). (Scale bar: 25µm) (C) FRET efficiencies calculated from CLSM images with FRET SE method (n=30). Green fluorescence represents the fluorescence of DiO. Red fluorescence represents the fluorescence of DiI.

Taken together with the cellular uptake study in Figure 3, the current results possibly indicated that the internalization speed of miktoarm copolymers was slower than that of core-loaded hydrophobic probes. This probably resulted from the hyperbranched hydrophobic block PCL, which easily embedded into cell membranes but was hard to be transferred into cells. Another proposition was that the PEG shell of PEG_113_-(*hb*-PG)_15_-*g*- PCL_22_ miktoarm copolymer would facilitate the fast release of its core loaded hydrophobic probes as result of PEG bridging between the phospholipid membranes of its adjacent cells^38^. While MCNs released the core-loaded hydrophobic probes into cells, the probes were re-encapsulated into endocytic vesicles and transported into the cells.

### *In vivo* trafficking studies of FRET MCNs on MCF-7 tumor-bearing mice. *A) FRET measurements*

To monitor the dynamics of MCNs transferring *in vivo*, two near-infrared carbocyanine dyes DiD and DiR were loaded into MCNs. The chosen dyes DiD (donor: excitation [ex] /emission [em], 644/665 nm) and DiR (acceptor: excitation [ex] /emission [em], 750/780 nm) display a spectral overlap between the donor emission and the acceptor excitation spectrum^39^. Moreover, the near infrared excitation and emission wavelength of both DiD and DiR could effectively reduce the interference of animal auto-fluorescence in the following *in vivo* study^40^. Figure 4A shows the fluorospectra of MCNs loaded with DiD (DiD MCNs), MCNs loaded with DiR (DiR MCNs), MCNs loaded with DiD and DiR (DiD/DiR MCNs) and mixtures of DiD MCNs and DiR MCNs (DiD MCNs & DiR MCNs Mixtures). To certify the occurrence of FRET, all samples were excited at 644 nm. For DiD/DiR MCNs, a strong DiR signal was observed due to the close proximity of DiD and DiR in the core, with a FRET ratio of 1.25 (I_DiR_/I_DiD_). No FRET signal was observed in the sample of DiD MCNs & DiR MCNs Mixtures, with a FRET ratio of 0.12. Figure 4B shows *in vivo* NIR fluorescence imaging of these respective preparations. Upon excitation at 644 nm, strong emission of DiR FRET-mediated by DiD indicated compact encapsulation of DiD and DiR in same MCNs. In comparison, DiD MCNs & DiR MCNs Mixtures have only a slight emission in the FRET channel because DiD and DiR was not directly condensed with each other, but associated with MCNs respectively.

**Figure 4.**
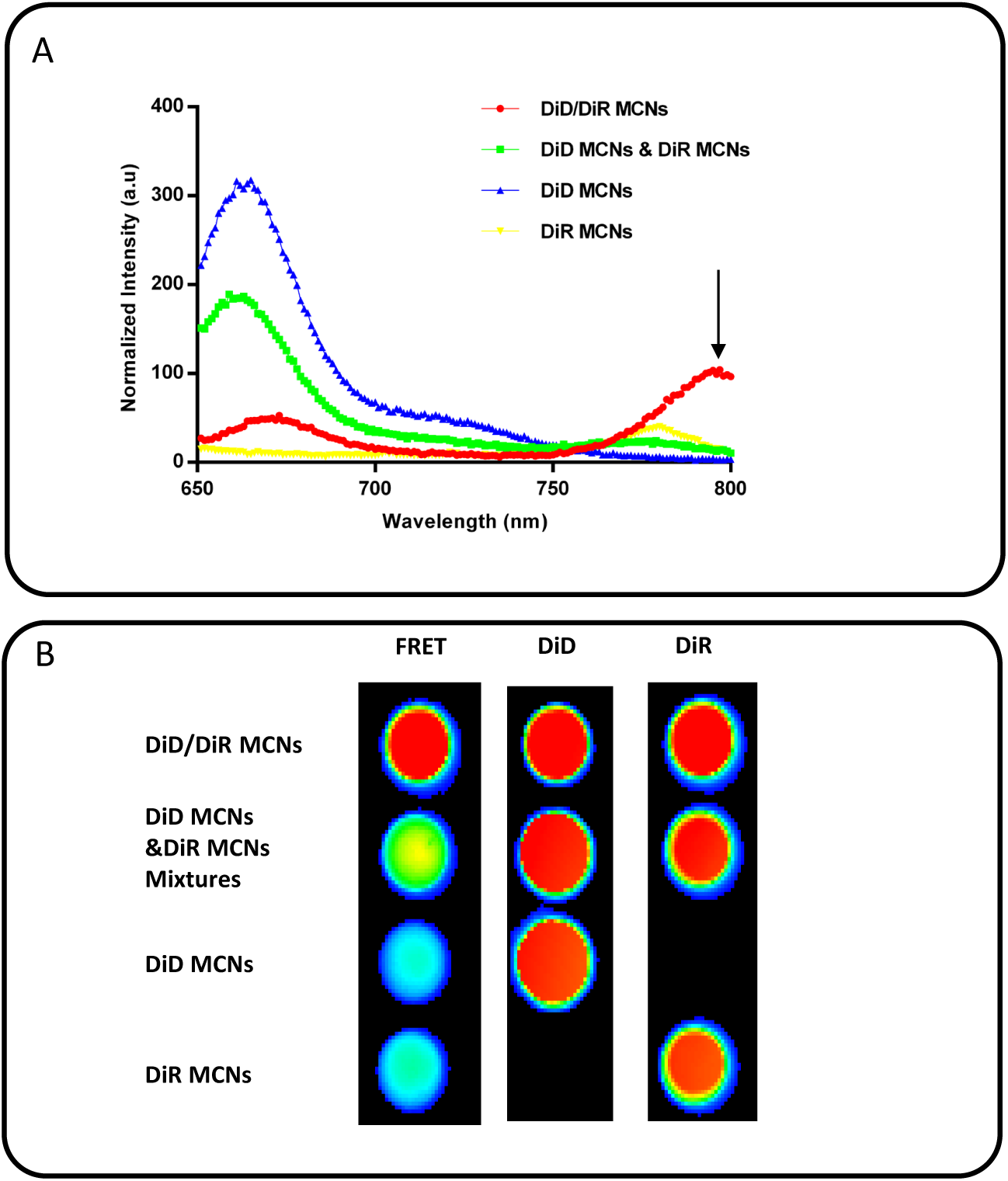
Fluorospectra and *in vivo* NIR fluorescence imaging of MCNs. (A) Fluorospectra of MCNs loaded with DiD (DiD MCNs), MCNs loaded with DiR (DiR MCNs), MCNs co-loaded with DiD and DiR (DiD/DiR MCNs), mixtures of MCNs loaded with DiD and MCNs loaded with DiR (DiD MCNs & DiR MCNs Mixtures). (B) *In vivo* NIR fluorescence imaging of of DiD/DiR MCNs and DiD MCNs & DiR MCNs Mixtures.

### B) In vivo fate of FRET MCNs on MCF-7 tumor-bearing mice

*In vivo* fate of DiD/DiR MCNs and DiD MCNs & DiR MCNs Mixtures was studied on MCF-7 xenografts in nude mice by NIR fluorescence imaging. As shown in Figure 5A, from 1 h to 8 h, the FRET- mediated DiR fluorescence intensity in DiD/DiR MCNs group was significantly higher than that of the DiD MCNs & DiR MCNs Mixtures group. Consistently, the DiD (donor) fluorescence intensity in DiD/DiR MCNs group was significantly lower than that of the DiD MCNs & DiR MCNs Mixtures group. This indicates the integrity of the DiD/DiR MCNs and the donor DiD transfers the excitation energy to the acceptor DiR due to the close proximity. From 8 h to 24 h, the fluorescence intensity of DiD and that of FRET-mediated DiR was almost the same in both DiD/DiR MCNs and DiD MCNs & DiR MCNs Mixtures groups. The FRET-mediated DiR fluorescence intensity achieved its highest level at 24 h. This observation demonstrated that in the first or tissue phase (1-8 h), our constructed MCNs accumulated, but were not internalized by tumor tissue through the EPR effect. As a result, fluorescence intensity of FRET- mediated DiR in the DiD/DiR MCNs group was higher than that of the DiD MCNs & DiR MCNs Mixtures group. In the second or cellular phase (8-24 h), we can infer the following process from the observation. 1) Similar to the first phase, more MCNs slowly accumulated in tumor tissue and ultimately reached the highest level at 24 h. 2) More importantly, MCNs were gradually internalized into MCF-7 tumor cells in this phase. Consistent with the result from trafficking studies *in vitro* (Figure 3A), fluorescent probes in MCNs were most likely concentrated in endocytic vesicles in the process of cellular internalization and intracellular sorting. FRET will take place in the concentrated DiD/DiR endocytic vesicles, same as FRET observed in DiD/DiR MCNs. Consequently, in the case of DiD MCNs & DiR MCNs Mixtures, the FRET-mediated DiR intensity gradually reached the same level with that of the DiD/DiR MCNs group in the second phase.

**Figure 5.**
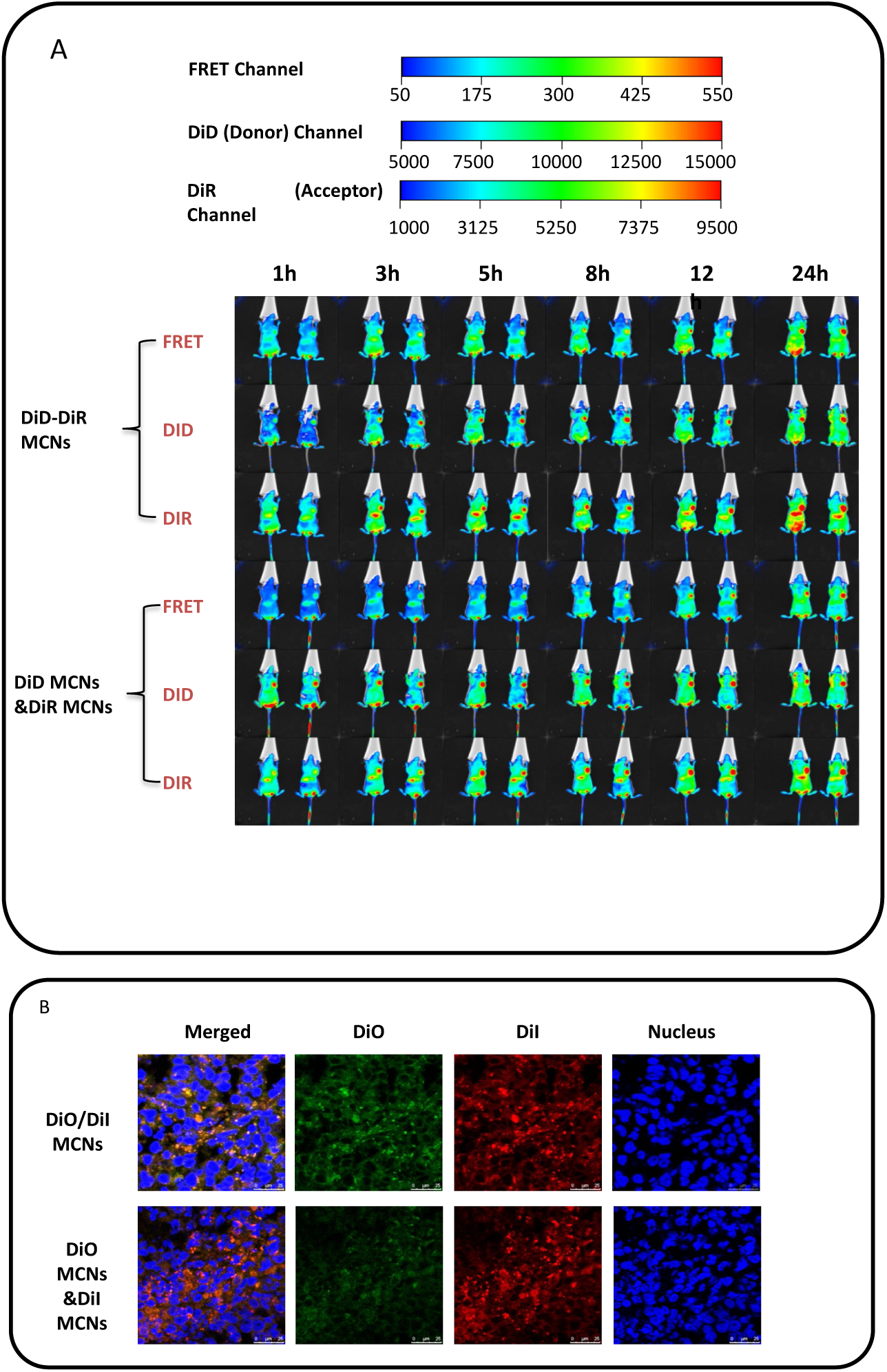
*In vivo* fate of FRET MCNs on MCF-7 tumor-bearing mice. (A) NIR fluorescence imaging of MCF-7 tumor-bearing mice at 1, 3, 5, 8, 12 and 24 h after iv injection of DiD/DiR MCNs and DiD MCNs & DiR MCNs Mixtures. (B) CLSM images of fluorescence detection of DiO and DiI in tumor cryosections after iv injection of DiD/DiR MCNs and DiD MCNs & DiR MCNs Mixtures.

In tumor tissues, the distribution of DiO and DiI in tumor cells were subsequently examined with CLSM after tail vein injection of either DiO/DiI MCNs or DiO MCNs & DiI MCNs Mixtures. The distribution accorded with the FRET imaging and analysis by NIR fluorescence imaging (Figure 5B). Collectively, the *in vitro* and *in vivo* trafficking studies confirmed the correlation of cellular endocytosis process of MCNs loaded with hydrophobic fluorescent probes.

### Cellular uptake of MCNs with different copolymer mass ratios

After the clarification of the MCN trafficking pathway *in vitro* and *in vivo*, the underlying interaction mechanism of MCNs with the cell membrane was assessed by comparing the uptake amount and speed of different MCNs. Four kinds of DiO/DiI MCNs were prepared and the mass ratio of copolymer to DiO/DiI probes was 500:1, 750:1, 1500:1 and 3000:1. Our results suggested that with the increase of mass ratio of copolymer, the uptake of DiO and DiI significantly decreased (Figure 6). Similar observations were noticed in both flow cytometry (Figure 6A, 6B) analysis and CLSM studies (Figure 6C). This result suggested that the mass ratio of the copolymer affected internalization of the hydrophobic fluorescence probes.

**Figure 6.**
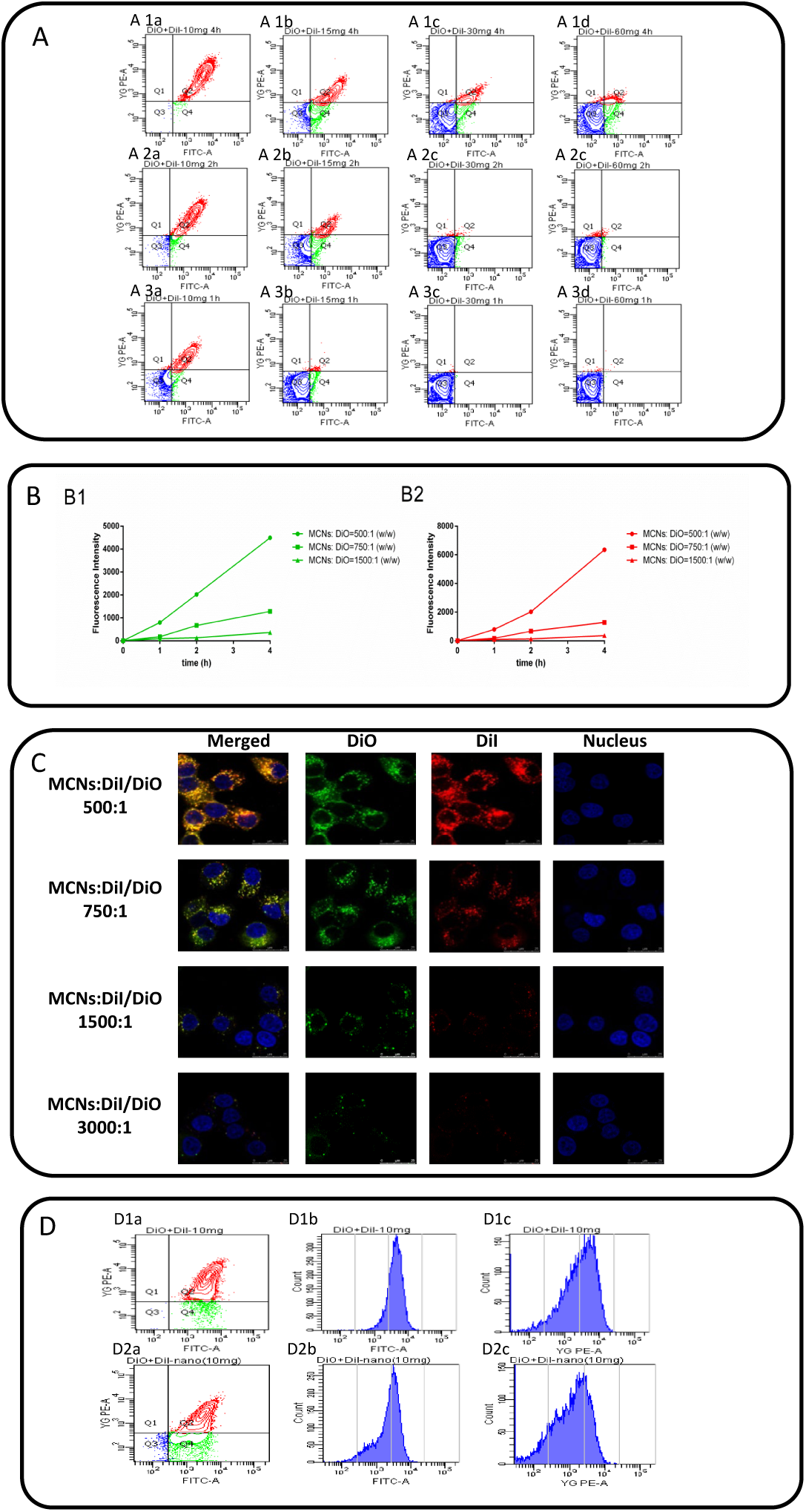
Cellular uptake of MCNs with different copolymer ratio. (A) Fluorescence intensity of MCF-7 cells analyzed by flow cytometric after treated with DiO/DiI MCNs at different copolymer ratios to probes: 500:1 (Aa), 750:1 (Ab), 1500:1 (Ac), 3000:1(Ad) at 1(A1), 2 (A2), 4h (A3). Green represents the fluorescence of DiO. Red represents the fluorescence of DiI. (B) Uptake speed of DiO (B1) and DiI (B2) at different copolymer ratios to probes: 500:1, 750:1 1500:1. (C) CLSM images of fluorescence detection of DiO and DiI in MCF-7 cells with DiO/DiI MCNs at different copolymer ratios to probes: 500:1, 750:1, 1500:1, 3000:1 at 4 h. (D) Flow cytometry analysis for MCNs loaded with DiO (Db) and DiI (Dc) without preincubation (D1) or with preincubation (D2) of blank MCNs. The concentration of DiO was 2 μg/mL, the concentration of DiI was 2 μg/mL for all formulations that has DiO or DiI.

To further investigate factors affecting cellular uptake, we calculated the uptake speed of three kinds of DiO/DiI MCNs (the uptake of 3000:1 mass ratio of copolymer to DiO/DiI probes MCNs was too insignificant to be calculated and thus wasn’t included in further studies). The fluorescence intensity of DiO or DiI in each group was plotted with time points. As indicated in Figure 6B, three groups all showed burst-free uptake in a linearly increasing. This suggested that different MCNs interacted with cells possibly via the same endocytosis pathway.

Moreover, in order to further assess the influence of copolymers, we pre-incubated MCF-7 cells with blank MCNs (MCNs without loading fluorescent probes) for 1 h, copiously washed the cells with PBS, and incubated them with DiO/DiI MCNs (the mass ratio of copolymer to DiO/DiI probes is 500:1) for another 2 h. It was found that the uptake of DiO and DiI decreased after pre-incubation with blank MCNs (Figure 6D), presumably suggesting that the amphiphilic hyperbranched copolymers might embed into the cell membrane after preincubation, which affected the fluidity of the cell membrane and therefore hindered the subsequent cellular uptake of DiO/DiI MCNs.

### *In vivo* distribution of DiR MCNs with different copolymer ratios

To develop a correlation between *in vitro* cellular uptake and *in vivo* distribution of MCNs with different copolymer ratios, the tumor distribution of DiR MCNs with different copolymer ratios was investigated by near-infrared fluorescence (NIRF) imaging. Figure 7A shows the *in vivo* NIRF images of MCF-7 tumor-bearing mice at different time points after IV injection of DiR MCNs with different copolymer ratios (the loaded DiR concentration in all groups was 10 µg/mL). After the injection of different MCNs, a strong fluorescence of DiR in the tumor region was observed in all groups (Figure 7B) and achieved the strongest fluorescence intensity at 24 h. Most interestingly while maintaining the same amount of DiR, the highest copolymer ratio group exhibited the highest and longest tumor targeting. A recent study showed that DiR existed in a nonquenched state in the cores of polymeric micelles^40^. Moreover, with the increase of copolymer ratio, it was more difficult for DiR to leak out from MCNs. As a result, the group with the highest copolymer ratio showed the most localization in the tumor region via the EPR effect. Furthermore, consistent with the *in vitro* cellular uptake assay, the results also demonstrated that, with the increase of copolymer ratio, the clearance rate decreased. After 96 h of injection, DiR MCNs were still found in the peripheral region of tumor in 3000:1 (MCNs: DiR) group. Increased copolymer ratio created a stronger stealth system, which prevented the tumor cell uptake of DiR and thus slowed down the clearance rate. In another perspective, our results also suggested that the polymer and dye ratio should be optimized when using this delivery system to provide a basis for enhancing surgical guidance via NIR visualization of tumors. Although a higher polymer ratio would slow down the clearance rate, it would also amplify the tumor region at the same time due to their slow internalization and peripheral tumor distribution.

**Figure 7.**
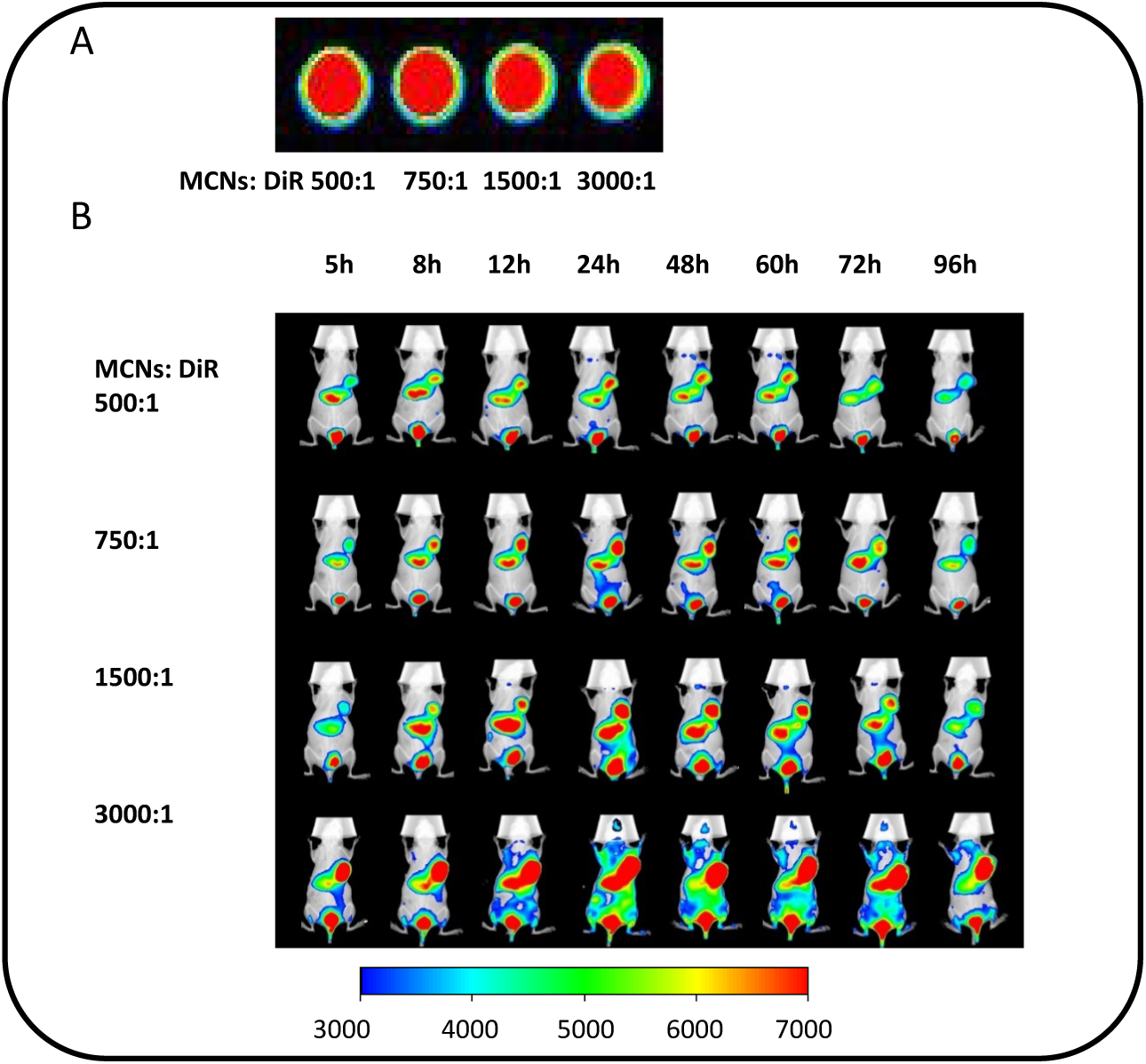
In vivo NIR fluorescence imaging of MCF-7 tumor bearing mice at the indicated time point after iv injection of DiR MCNs with different ratio of copolymers. (A) NIR fluorescence imaging of DiR MCNs with different ratio of copolymers. (B) In vivo real-time imaging of MCF-7 tumor bearing mice. The concentration of DiR was 10 μg/mL for all formulations at a dose of 100 μg/kg.

## Conclusion

Understanding the underlying endocytosis mechanism of miktoarm copolymer micelles is critical for the design of non-linear copolymers as effective drug carriers. Despite numerous studies examining the endocytosis and intracellular trafficking of polymeric micelles^26, 38, 41^, whether the endocytosis mechanism discovered *in vitro* can be applied *in vivo* remains unknown. This study is essential considering that, only by confirming the *in vitro*-*in vivo* correlation of copolymer nanomicelles in cellular uptake and trafficking could we then validate and apply the mechanism discovered in the cellular level to design new copolymers for drug delivery. Our findings demonstrated that the *in vitro* and *in vivo* mechanism of cellular uptake and intracellular trafficking of MCNs involved a clear and similar sequence of events: interaction of MCNs with the cell membrane induced hyperbranched block PCL embedding into the cytomembrane, which resulted in the release of loaded hydrophobic fluorescence probes from the cores of MCNs and was followed by time-dependent intracellular clustering within endocytic vesicles. Moreover, the uptake amount and speed of loaded fluorescent probes were correlated with the mass ratios of MCNs *in vitro* and *in vivo*. A high mass ratio of MCNs to loaded hydrophobic probes also resulted in slower clearance rate of loaded fluorescent probes in the peripheral region of the tumor. This study not only highlights the need for more thorough studies regarding the *in vitro-in vivo* transport mechanism correlation of polymeric nanomaterials but also provides a universal method of visualizing this relationship both *in vitro* and *in vivo*. Continued research in this topic would help the design of more effective nanomaterials platforms for biomedical applications. The current work initiates the impetus to synthesize more kinds of hyperbranched copolymers to navigate the characteristics of the cell membrane for better cellular targeting and penetrating.

## Figure legend

**Table S1.** Characteristics of PEG_113_-(*hb*-PG)_15_-*g*-PCL_22_ miktoarm copolymer nanomicelles

| Preparations | Particle Size (nm) | Polydispersity | Zeta potential(mV) | Encapsulation Efficiency(%) |
| --- | --- | --- | --- | --- |
| MCNs | 45.17 ± 2.3 | 0.228±0.01 | -0.013±0.02 | - |
| DiO MCNs | 51.41 ± 4.1 | 0.16±0.03 | -0.787±0.11 | 90.6%±0.5 |
| DiI MCNs | 52.46±1.2 | 0.152±0.02 | -1.78±0.25 | 92.1%±1.2 |
| DiD MCNs | 55.20±3.5 | 0.193±0.02 | -0.925±0.11 | 93.4%±1.0 |
| DiR MCNs | 53.31±3.1 | 0.219±0.03 | -1.347±0.21 | 92.4%±0.9 |

**Figure S1.**
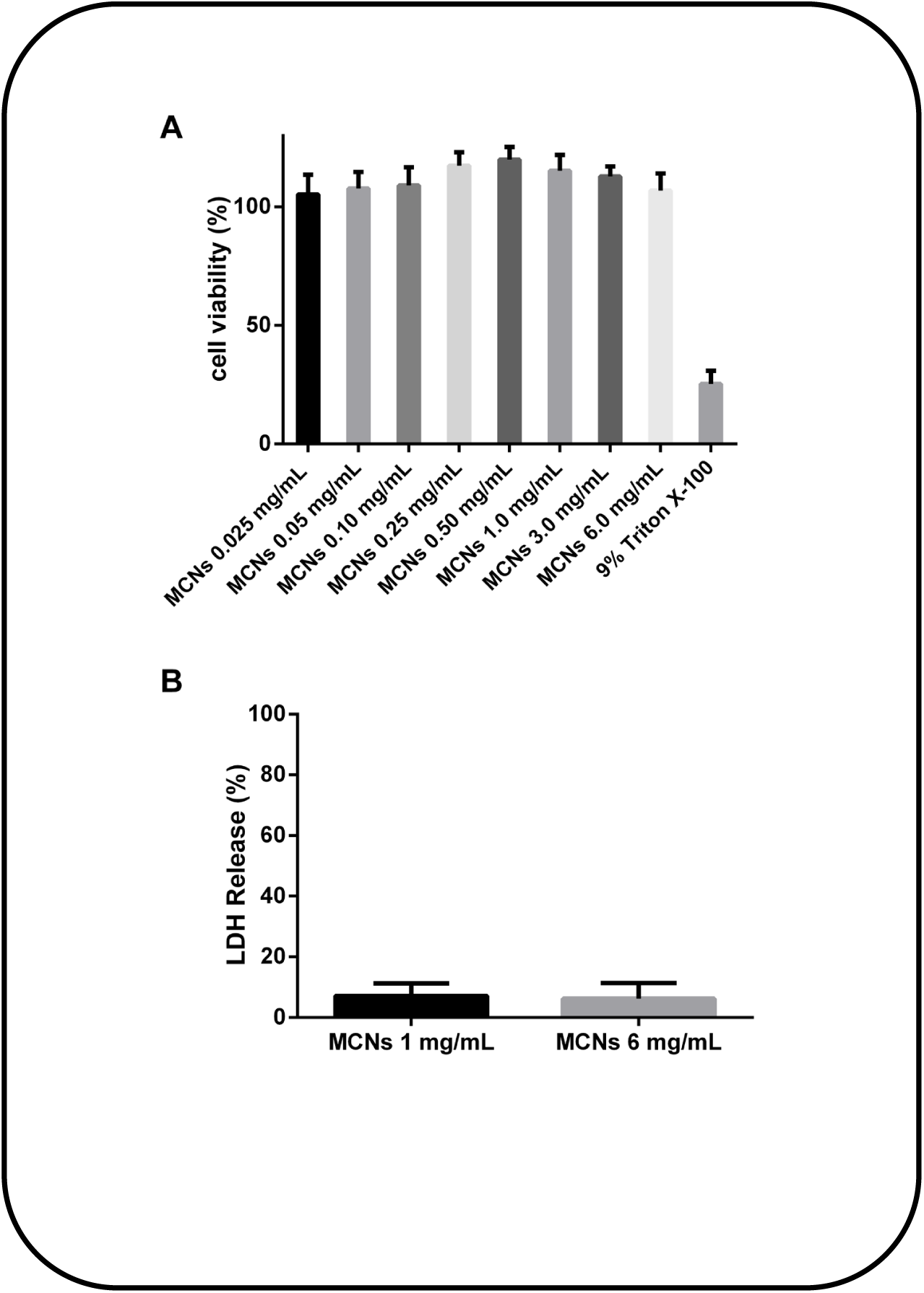
Results of the cell viability after being treated with MCNs with different concentration, determined by (A) the SRB assay (n=5) and (B) the LDH assay (n=3).

**Figure S2.**
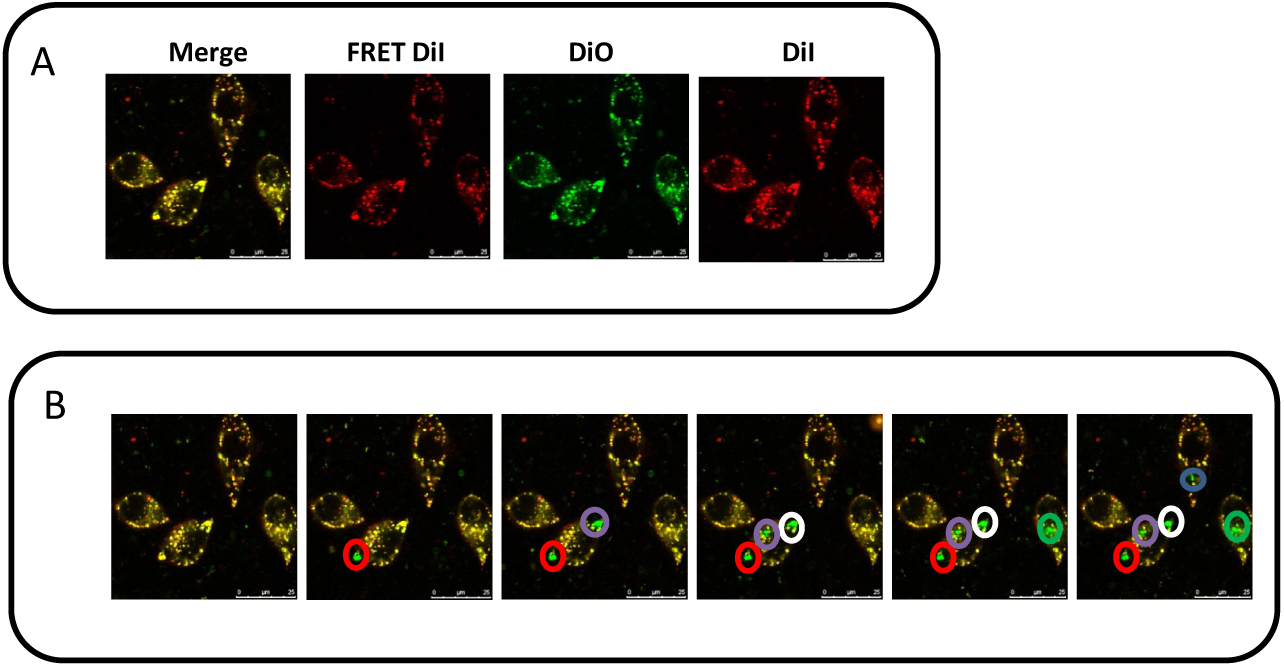
CLSM image of MCF-7 cells incubated with DiO/DiI MCNs for 4 h. (A) Images obtained to calculate FRET efficiencies by FRET Sensitized Emission (SE) method. (B) Images obtained to calculate FRET efficiencies by FRET Acceptor PhotoBleaching (AB) method. Circled region was photobleaching points. Five points were selected to be bleached accordingly. Green fluorescence represents the fluorescence of DiO. Red fluorescence represents the fluorescence of DiI. (Scale bar: 25 µm)

## Author contributions

The manuscript was written through contributions of all authors. All authors have given approval to the final version of the manuscript.

## Notes

The authors declare no competing financial interest.

## Acknowledgment

This work was supported by the Fundamental Research Funds for the National Natural Science Foundation of China (81821004 and 81690264 to Q. Zhang).

